# Developmental factors associated with decline in grip strength from midlife to old age: a British birth cohort study

**DOI:** 10.1101/327247

**Authors:** D Kuh, R Hardy, JM Blodgett, R Copper

## Abstract

Maintenance of muscle strength is important for healthy ageing, protecting against chronic disease and enabling independent living. We tested whether developmental factors were associated with grip strength trajectories between 53 and 69 years, and operated independently or on the same pathway/s as adult factors, in 3058 participants from a British birth cohort. Grip strength (kg) at ages 53, 60-64 and 69, was analysed using multilevel models, testing for age and sex interactions, to estimate associations with developmental factors (birthweight, growth parameters, motor and cognitive development) and childhood socioeconomic position (SEP) and investigate potential adult mediators. Heavier birthweight, beginning to walk on time, later puberty and greater weight 0-26 years in men, and earlier age at first standing in women, were associated with stronger grip but not with its decline; these associations were independent of adult factors. The slower decline in grip strength (by 0.068kg/year, 95% confidence interval (CI) 0.024,0.11 per 1SD, p=.003) in men with higher childhood cognition was attenuated by adult verbal memory which became increasingly positively associated with grip strength at older ages. Thus grip strength may increasingly reflect neural ageing processes. Targeting developmental factors to promote muscle development should increase the chance of independence in old age.

---

Maintaining musculoskeletal function for the maximal period of time, and preventing musculoskeletal disorders are important aspects of healthy ageing, enabling people to remain active and independent for longer.(1, 2) Worldwide, musculoskeletal disorders rank second in years lived with disability and fourth in disability-adjusted life years.(2) The role of muscle mass, strength and metabolic function is recognised for these disorders, and is becoming more widely appreciated in cardiovascular and other chronic diseases.(3, 4)

Hand grip strength is a commonly used indicator of muscle strength;(5) and an overall biomarker of ageing.(6–8) Average levels rise to a peak during the early 30s, plateau and then decline.(9–11) Weaker grip is associated with future morbidity, disability and mortality across populations of different ages, ethnicities and income levels,(4, 12–18) as is decline in grip strength.(19, 20) Adult risk factors, including height and adiposity, health conditions, cognition and health behaviours, have been associated with subsequent grip strength,(21, 22) and with age-related decline.(23–34) Developmental factors, such as birthweight, physical growth, motor and cognitive development, and childhood socioeconomic conditions are also related to adult grip strength.(35, 36) but evidence on whether they are associated with age-related decline is limited.(34, 37, 38)

A research gap is to understand whether developmental factors operate on the rate of decline in grip strength independently or on the same pathway/s as these adult factors to inform the timing and types of interventions that may modify this decline. Using three repeat measures of grip strength ascertained from age 53 to 69 years in a British birth cohort, we tested two hypotheses: (1) that higher birthweight, greater gains in height and weight during childhood and adolescence, and earlier puberty are associated with a greater grip strength but not its decline; and (2) that achieving motor milestones early or around the modal age, and higher childhood cognitive ability and SEP, are associated with greater grip strength and a slower rate of decline. Further, we expected that any associations between indicators of physical growth and grip strength to be mediated by adult health conditions and body mass index (BMI), and that any associations between motor and cognitive development, childhood SEP and grip strength to be mediated by education and adult cognition.

## METHODS

### Sample

The Medical Research Council National Survey of Health and Development (NSHD) is a sample of all births in one week in March 1946 in mainland Britain comprising 5,362 (2,547 female) individuals followed up 24 times, so far to age 69.(39) The maximum sample for these analyses comprised 3,058 participants with at least one measure of grip strength at ages 53, 60-64 or 69 years and known adult height and birthweight. Of the remaining 2,304 in the original birth cohort, 738 had died, 542 were living abroad, 270 had been lost to follow-up, and 166 had not provided all necessary data. Ethical approval for the most recent visit was given by Queen Square Research Ethics Committee (13/LO/1073) and Scotland A Research Ethics Committee (14/SS/1009). Written informed consent was provided by participants for each visit.

### Grip strength

During nurse assessments at ages 53 and 60-64, grip strength was measured in kilograms isometrically using a Nottingham electronic handgrip dynamometer; during a nurse home visit at age 69, a Jamar Plus+ Digital Hand dynamometer was used. A randomised repeated-measurements cross-over trial found no statistically significant differences in values when comparing these two devices.(40) We applied the same standardised protocols and used the maximum of the first four measures (two in each hand) at each age.

### Childhood factors

*Birthweight:* Birthweight, extracted from birth records to the nearest quarter pound, was converted to kilograms.

*Physical growth:* The SITAR model of growth curve analysis was used to estimate individual patterns of growth in height and weight between 0 and 26 years.(41, 42) Heights and weights were measured using standardised protocols at ages 2, 4, 6, 7, 11 and 15, and self-reported at ages 20 and 26. The NSHD data were augmented by height and weight data between 5 and 19 years from the ALSPAC cohort to provide additional information at intermediate ages. Subject-specific random effects were obtained for size, tempo and velocity (in SD units) for height and weight.(41) Later puberty is indicated by positive tempo values and earlier puberty by negative values. (41)

*Motor and cognitive development:* Age (in months) at first sitting, standing and walking was based on maternal recall at age 2. At age 15, a standardised measure of childhood cognitive ability was derived from the Heim AH4 test of fluid intelligence, the Watts Vernon reading test and a test of mathematical ability.(43, 44) Standardised scores from similar tests at ages 11 or 8 were used if missing at age 15 as participants maintained similar ranking across time.

*Childhood SEP:* Father’s occupation at age 4 (or at age 11 or age 15 if missing at age 4), based on the Registrar General’s Social Classification, distinguished three groups: high (I or II), middle (IINM or IIM) and low (IV and V).

### Adult factors

*Height and adiposity*: At ages 53, 60-64 and 69, height (cm) and weight (kg) were measured using standard protocols and BMI (kg/m^2^) was calculated; standardised scores were used in analyses.

*Health conditions:* At age 53, a summary of health conditions was a count (0-4) of the presence of knee osteoarthritis, hand osteoarthritis (both based on clinical assessment), severe respiratory symptoms and other potentially disabling or life threatening conditions.(25)

*Education and verbal memory*: Highest educational attainment by age 26 distinguished those with a degree or higher, advanced secondary, ordinary secondary, other or no qualifications. At ages 53, 60-64 and 69, verbal memory was assessed using a 15 item word list task over three trials (range 0-45),(45) and was converted to a standardised score.

*Other adult covariates:* At ages 53, 60-64 and 69, participants reported if they smoked and how many times they had taken part in any sports or vigorous leisure activities in the last month (grouped into more than 5 times a month, 1-4 times a month or not at all). Adult SEP, assessed by own occupation at age 53 (or at earlier ages if missing at age 53), distinguished the same three groups as for father’s occupation.

### Statistical analysis

Stata v14.2 was used for all analyses. We fitted multilevel models which account for the correlation of repeated measures of grip strength within individuals. Preliminary multilevel models tested whether adult height was associated with grip strength and remained constant with age,(33) and whether men had stronger grip but a faster rate of decline than women, as expected.(23, 28, 29, 33, 37, 46, 47)

In the main analysis, models were run separately for men and women because of evidence of sex interactions with age and other covariates; these are reported where statistically significant. All models were adjusted for height, with change in grip strength modelled by a linear age term, and with the intercept and slope fitted as random effects. We also tested for age interactions with each of the risk factors, including those significant at the 0.1 level in models.

To investigate developmental risk factors and grip strength, we first investigated separately the associations with birthweight, physical growth, motor development, cognitive development and childhood SEP. For physical growth, we included all SITAR parameters in the same multilevel models. For motor development, we ran models for age at first sitting, standing and walking, first separately and then mutually adjusted.

Then we took the developmental factors associated with grip strength at the end of the first stage of the analysis and adjusted in turn for each group of adult factors, having shown in supplementary analyses how they were associated with grip strength.

In sensitivity analyses, to assess potential bias introduced by: (a) excluding those participants with no valid observations who were unable to perform the test for health reasons, and (b) mortality or other attrition during follow-up, we reran the multilevel models (1) giving a value representing the midpoint of the lowest sex-specific fifth of grip strength to participants unable for health reasons (n=29, 81 observations); and (2) including binary indicators for mortality (n=287) and attrition (n=601).(27, 48)

## RESULTS

### Characteristics of the sample and preliminary analyses

Mean levels of grip strength, birthweight, and adult height were greater in men than women; women had more health conditions and lower SEP than men (Table 1). Mean grip strength declined between ages 53 and 69 by 7.5kg for men and 3.6kg for women; thus the difference between men and women attenuated over time, although the mean sex difference remained substantial at age 69 (16.1kg) (Table 1). Preliminary multilevel models confirmed that adult height was strongly associated with grip strength (3.2 kg, 95% confidence interval (CI) 2.8,3.5 per 1SD increase in height, p=<.001) and remained constant with age, and that men had stronger grip and a faster rate of decline than women (p-value<.001 for sex interaction with height).

**Table 1.**
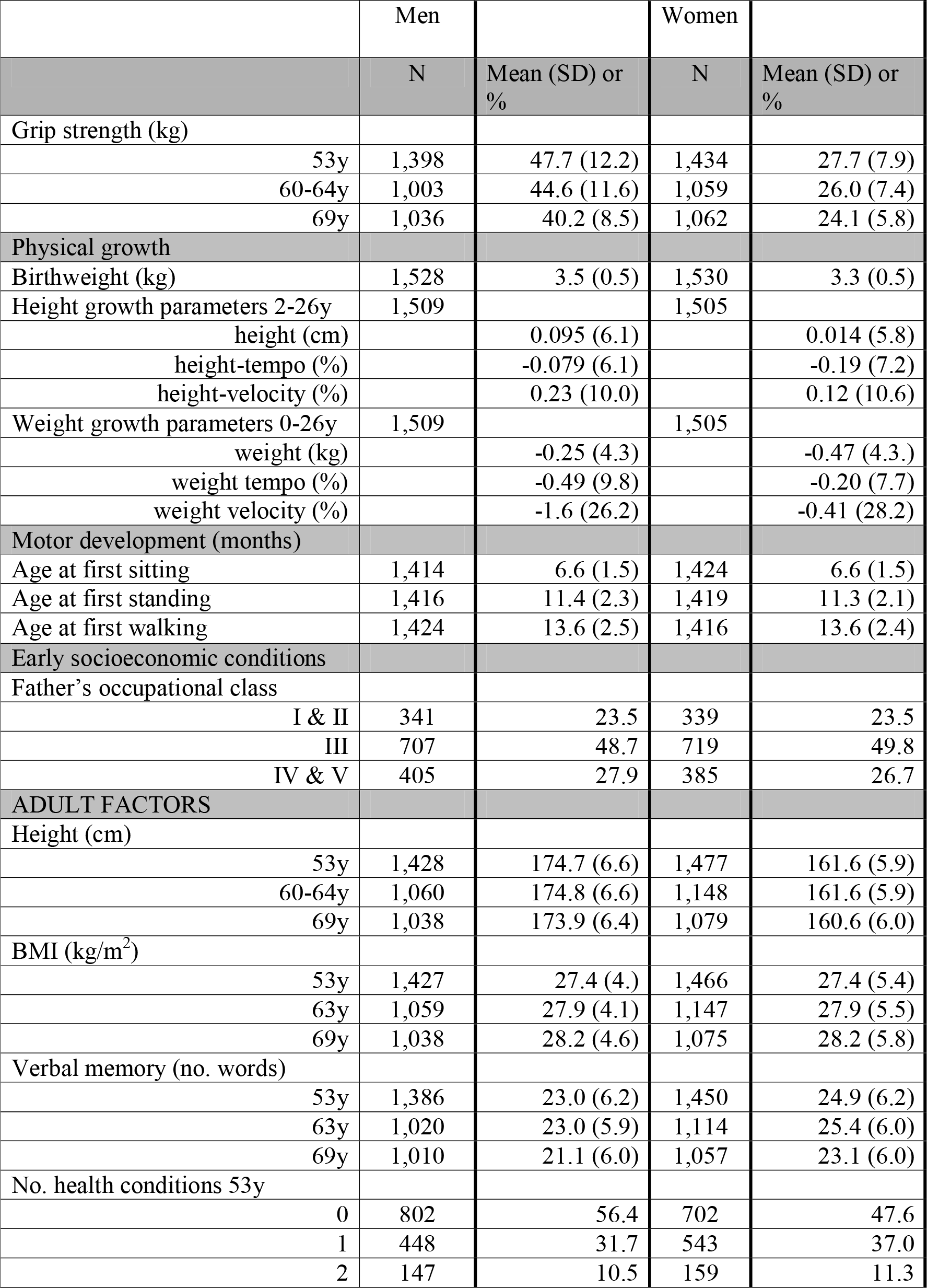
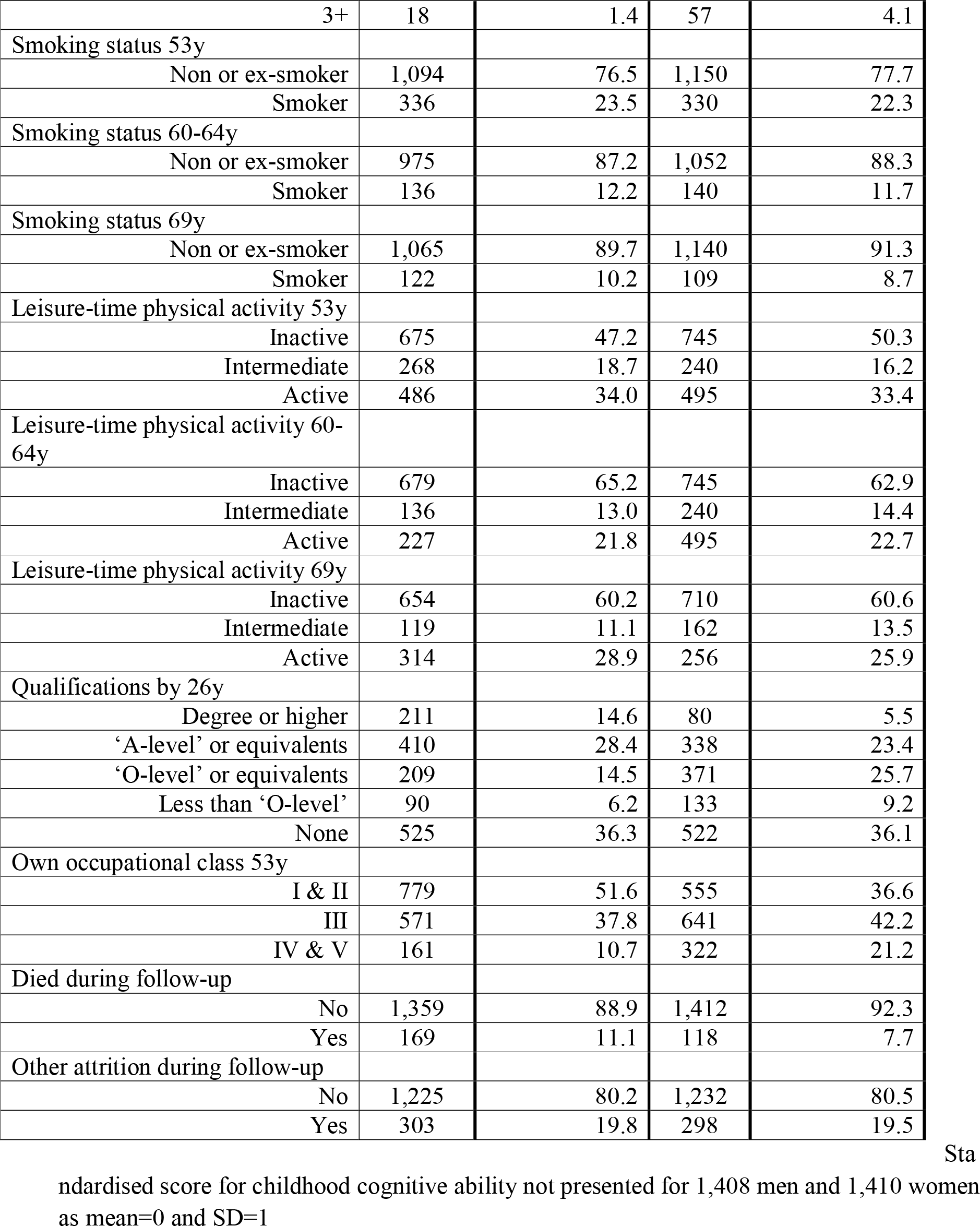
Characteristics of the Sample of 1,528 Men and 1,530 Women in the MRC National Survey of Health and Development With at Least One Measure of Grip Strength at ages 53, 60-64, or 69 and known Height and Birthweight

|  | Men |  | Women |  |
| --- | --- | --- | --- | --- |
|  | N | Mean (SD) or % | N | Mean (SD) or % |
| Grip strength (kg) |  |  |  |  |
| 53y | 1,398 | 47.7 (12.2) | 1,434 | 27.7 (7.9) |
| 60-64y | 1,003 | 44.6 (11.6) | 1,059 | 26.0 (7.4) |
| 69y | 1,036 | 40.2 (8.5) | 1,062 | 24.1 (5.8) |
| Physical growth |  |  |  |  |
| Birthweight (kg) | 1,528 | 3.5 (0.5) | 1,530 | 3.3 (0.5) |
| Height growth parameters 2-26y | 1,509 |  | 1,505 |  |
| height (cm) |  | 0.095 (6.1) |  | 0.014 (5.8) |
| height-tempo (%) |  | -0.079 (6.1) |  | -0.19 (7.2) |
| height-velocity (%) |  | 0.23 (10.0) |  | 0.12 (10.6) |
| Weight growth parameters 0-26y | 1,509 |  | 1,505 |  |
| weight (kg) |  | -0.25 (4.3) |  | -0.47 (4.3.) |
| weight tempo (%) |  | -0.49 (9.8) |  | -0.20 (7.7) |
| weight velocity (%) |  | -1.6 (26.2) |  | -0.41 (28.2) |
| Motor development (months) |  |  |  |  |
| Age at first sitting | 1,414 | 6.6 (1.5) | 1,424 | 6.6 (1.5) |
| Age at first standing | 1,416 | 11.4 (2.3) | 1,419 | 11.3 (2.1) |
| Age at first walking | 1,424 | 13.6 (2.5) | 1,416 | 13.6 (2.4) |
| Early socioeconomic conditions |  |  |  |  |
| Father's occupational class |  |  |  |  |
| I & II | 341 | 23.5 | 339 | 23.5 |
| III | 707 | 48.7 | 719 | 49.8 |
| IV & V | 405 | 27.9 | 385 | 26.7 |
| ADULT FACTORS |  |  |  |  |
| Height (cm) |  |  |  |  |
| 53y | 1,428 | 174.7 (6.6) | 1,477 | 161.6 (5.9) |
| 60-64y | 1,060 | 174.8 (6.6) | 1,148 | 161.6 (5.9) |
| 69y | 1,038 | 173.9 (6.4) | 1,079 | 160.6 (6.0) |
| BMI (kg/m<sup>2</sup>) |  |  |  |  |
| 53y | 1,427 | 27.4 (4.) | 1,466 | 27.4 (5.4) |
| 63y | 1,059 | 27.9 (4.1) | 1,147 | 27.9 (5.5) |
| 69y | 1,038 | 28.2 (4.6) | 1,075 | 28.2 (5.8) |
| Verbal memory (no. words) |  |  |  |  |
| 53y | 1,386 | 23.0 (6.2) | 1,450 | 24.9 (6.2) |
| 63y | 1,020 | 23.0 (5.9) | 1,114 | 25.4 (6.0) |
| 69y | 1,010 | 21.1 (6.0) | 1,057 | 23.1 (6.0) |
| No. health conditions 53y |  |  |  |  |
| 0 | 802 | 56.4 | 702 | 47.6 |
| 1 | 448 | 31.7 | 543 | 37.0 |
| 2 | 147 | 10.5 | 159 | 11.3 |
| 3+ | 18 | 1.4 | 57 | 4.1 |
| Smoking status 53y |  |  |  |  |
| Non or ex-smoker | 1,094 | 76.5 | 1,150 | 77.7 |
| Smoker | 336 | 23.5 | 330 | 22.3 |
| Smoking status 60-64y |  |  |  |  |
| Non or ex-smoker | 975 | 87.2 | 1,052 | 88.3 |
| Smoker | 136 | 12.2 | 140 | 11.7 |
| Smoking status 69y |  |  |  |  |
| Non or ex-smoker | 1,065 | 89.7 | 1,140 | 91.3 |
| Smoker | 122 | 10.2 | 109 | 8.7 |
| Leisure-time physical activity 53y |  |  |  |  |
| Inactive | 675 | 47.2 | 745 | 50.3 |
| Intermediate | 268 | 18.7 | 240 | 16.2 |
| Active | 486 | 34.0 | 495 | 33.4 |
| Leisure-time physical activity 60-64y |  |  |  |  |
| Inactive | 679 | 65.2 | 745 | 62.9 |
| Intermediate | 136 | 13.0 | 240 | 14.4 |
| Active | 227 | 21.8 | 495 | 22.7 |
| Leisure-time physical activity 69y |  |  |  |  |
| Inactive | 654 | 60.2 | 710 | 60.6 |
| Intermediate | 119 | 11.1 | 162 | 13.5 |
| Active | 314 | 28.9 | 256 | 25.9 |
| Qualifications by 26y |  |  |  |  |
| Degree or higher | 211 | 14.6 | 80 | 5.5 |
| 'A-level' or equivalents | 410 | 28.4 | 338 | 23.4 |
| 'O-level' or equivalents | 209 | 14.5 | 371 | 25.7 |
| Less than 'O-level' | 90 | 6.2 | 133 | 9.2 |
| None | 525 | 36.3 | 522 | 36.1 |
| Own occupational class 53y |  |  |  |  |
| I & II | 779 | 51.6 | 555 | 36.6 |
| III | 571 | 37.8 | 641 | 42.2 |
| IV & V | 161 | 10.7 | 322 | 21.2 |
| Died during follow-up |  |  |  |  |
| No | 1,359 | 88.9 | 1,412 | 92.3 |
| Yes | 169 | 11.1 | 118 | 7.7 |
| Other attrition during follow-up |  |  |  |  |
| No | 1,225 | 80.2 | 1,232 | 80.5 |
| Yes | 303 | 19.8 | 298 | 19.5 |

Sta
Standardised score for childhood cognitive ability not presented for 1,408 men and 1,410 women as mean=0 and SD=1

### Developmental risk factors: multilevel models

In models adjusted for adult height and age, there were positive associations between birthweight and grip strength which were stronger in men than women and remained constant with age (Table 2a). In men, having a greater weight between birth and 26 years, and a later puberty (as indicated by a positive coefficient for height tempo) was associated with stronger grip. In women, shorter height, greater weight and a slower weight velocity between birth and 26 years were associated with stronger grip (Table 2b).

**Table 2.**
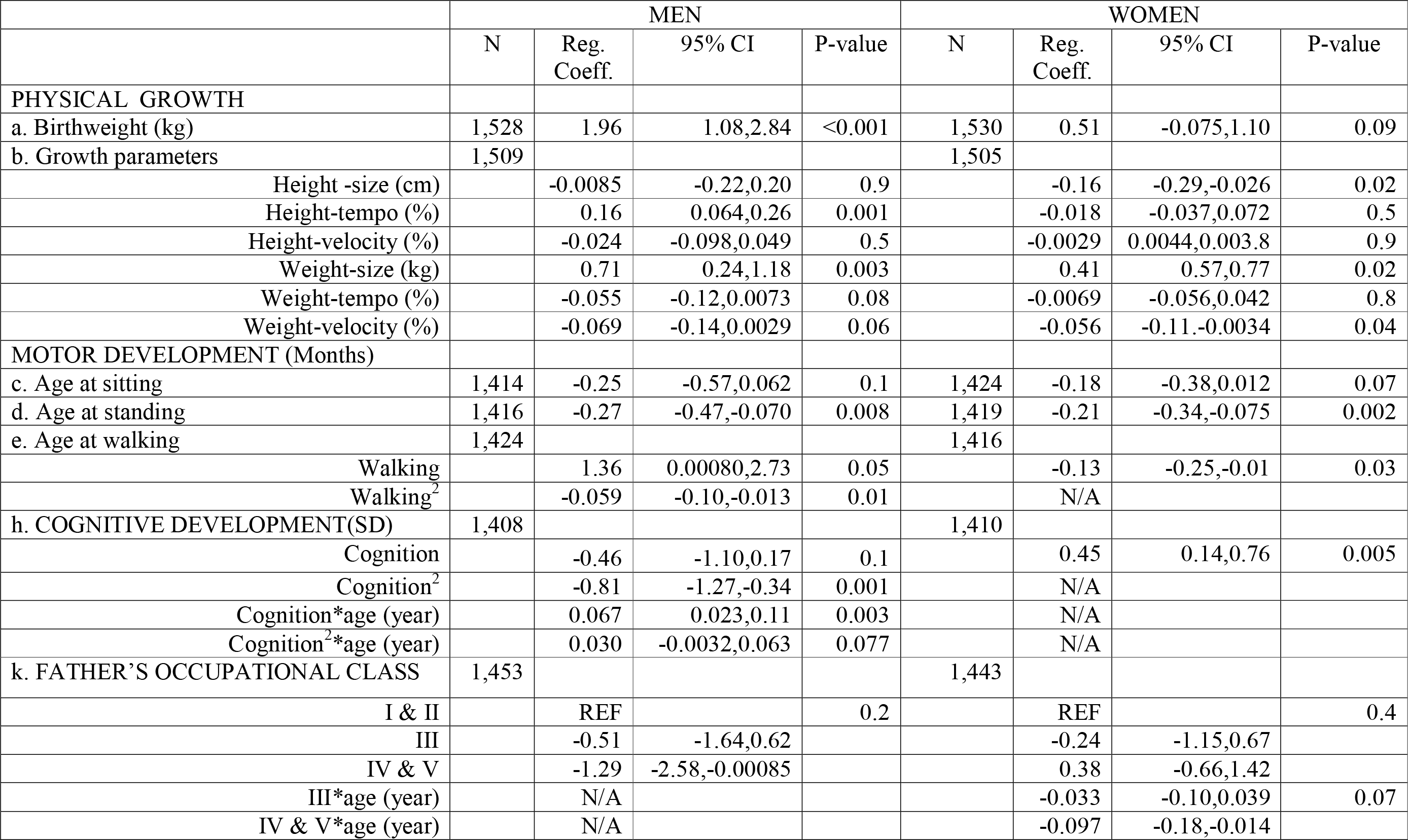
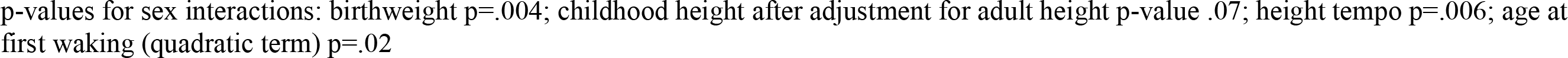
Estimates From Multilevel Models Showing Mean Differences in Grip Strength (kg) and Mean Differences in Grip Strength Changep-values for sex interactions: birthweight P=.004; childhood height after adjustment for adult height

|  | MEN |  |  |  | WOMEN |  |  |  |
| --- | --- | --- | --- | --- | --- | --- | --- | --- |
|  | N | Reg. Coeff. | 95% CI | P-value | N | Reg. Coeff. | 95% CI | P-value |
| **PHYSICAL GROWTH** |  |  |  |  |  |  |  |  |
| a. Birthweight (kg) | 1,528 | 1.96 | 1.08,2.84 | <0.001 | 1,530 | 0.51 | -0.075,1.10 | 0.09 |
| b. Growth parameters | 1,509 |  |  |  | 1,505 |  |  |  |
| Height -size (cm) |  | -0.0085 | -0.22,0.20 | 0.9 |  | -0.16 | -0.29,-0.026 | 0.02 |
| Height-tempo (%) |  | 0.16 | 0.064,0.26 | 0.001 |  | -0.018 | -0.037,0.072 | 0.5 |
| Height-velocity (%) |  | -0.024 | -0.098,0.049 | 0.5 |  | -0.0029 | 0.0044,0.003.8 | 0.9 |
| Weight-size (kg) |  | 0.71 | 0.24,1.18 | 0.003 |  | 0.41 | 0.57,0.77 | 0.02 |
| Weight-tempo (%) |  | -0.055 | -0.12,0.0073 | 0.08 |  | -0.0069 | -0.056,0.042 | 0.8 |
| Weight-velocity (%) |  | -0.069 | -0.14,0.0029 | 0.06 |  | -0.056 | -0.11,-0.0034 | 0.04 |
| **MOTOR DEVELOPMENT (Months)** |  |  |  |  |  |  |  |  |
| c. Age at sitting | 1,414 | -0.25 | -0.57,0.062 | 0.1 | 1,424 | -0.18 | -0.38,0.012 | 0.07 |
| d. Age at standing | 1,416 | -0.27 | -0.47,-0.070 | 0.008 | 1,419 | -0.21 | -0.34,-0.075 | 0.002 |
| e. Age at walking | 1,424 |  |  |  | 1,416 |  |  |  |
| Walking |  | 1.36 | 0.00080,2.73 | 0.05 |  | -0.13 | -0.25,-0.01 | 0.03 |
| Walking<sup>2</sup> |  | -0.059 | -0.10,-0.013 | 0.01 |  | N/A |  |  |
| h. COGNITIVE DEVELOPMENT(SD) | 1,408 |  |  |  | 1,410 |  |  |  |
| Cognition |  | -0.46 | -1.10,0.17 | 0.1 |  | 0.45 | 0.14,0.76 | 0.005 |
| Cognition<sup>2</sup> |  | -0.81 | -1.27,-0.34 | 0.001 |  | N/A |  |  |
| Cognition*age (year) |  | 0.067 | 0.023,0.11 | 0.003 |  | N/A |  |  |
| Cognition<sup>2</sup>*age (year) |  | 0.030 | -0.0032,0.063 | 0.077 |  | N/A |  |  |
| k. FATHER'S OCCUPATIONAL CLASS | 1,453 |  |  |  | 1,443 |  |  |  |
| I & II |  | REF |  | 0.2 |  | REF |  | 0.4 |
| III |  | -0.51 | -1.64,0.62 |  |  | -0.24 | -1.15,0.67 |  |
| IV & V |  | -1.29 | -2.58,-0.00085 |  |  | 0.38 | -0.66,1.42 |  |
| III*age (year) |  | N/A |  |  |  | -0.033 | -0.10,0.039 | 0.07 |
| IV & V*age (year) |  | N/A |  |  |  | -0.097 | -0.18,-0.014 |  |

p-values for sex interactions: birthweight $p=.004$ ; childhood height after adjustment for adult height $p\text{-value } .07$ ; height tempo $p=.006$ ; age at first waking (quadratic term) $p=.02$

Later ages of attaining infant motor milestones (for sitting, standing and walking) were associated with weaker grip which remained constant with age (Table 2c-e). The only exception was the inverse U-shaped relationship between age at first walking and grip strength in men. When all three motor milestones, together with height were included together, only the inverse U-shaped relationship between age at first walking and grip strength (for men) and the inverse association between age at first standing and grip strength (for women) remained (Supplementary Table 1); so only these variables were carried forward.

In men, there was an inverse U-shaped relationship between childhood cognitive ability and grip strength at age 53 (Table 2f). The age interaction terms show that the linear term strengthened (by 0.067kg/year, 95% confidence interval (CI) 0.024, 0.11 per 1SD, P=0.003) and the quadratic term became weaker with age, such that men of higher childhood cognition showed a slower decline in grip strength (Figure 1). In women, higher childhood cognitive ability was associated with stronger grip and this remained constant with age.

**Figure 1.**
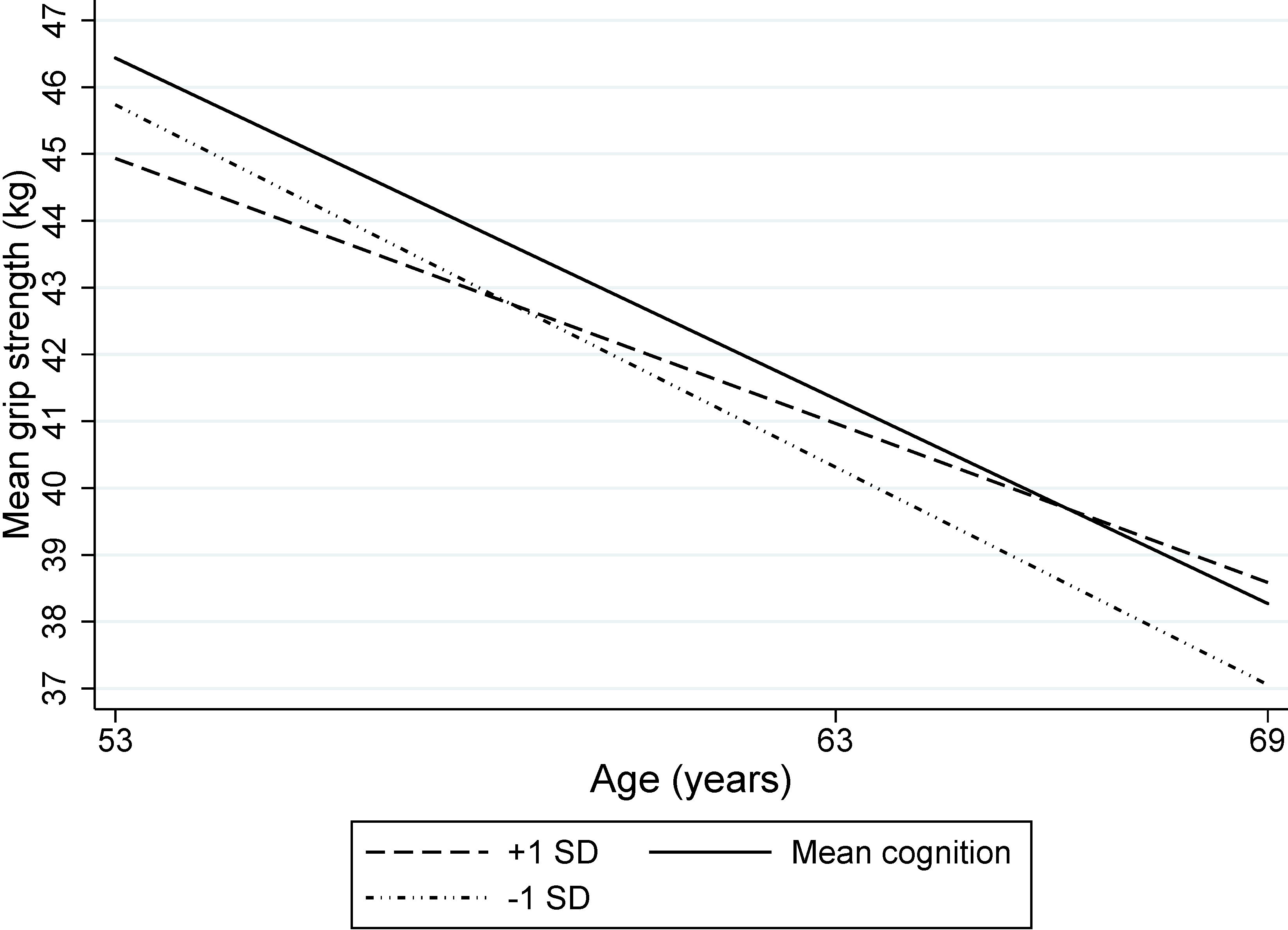
Mean grip strength (kg) by childhood cognition for men of mean height (based on 1408 men)

There was no association between childhood SEP and grip strength in men (Table 2g). However, in women there was weak evidence that the association grew stronger with age; women from social classes IV and V showed a faster decline in grip strength (Figure 2).

**Figure 2.**
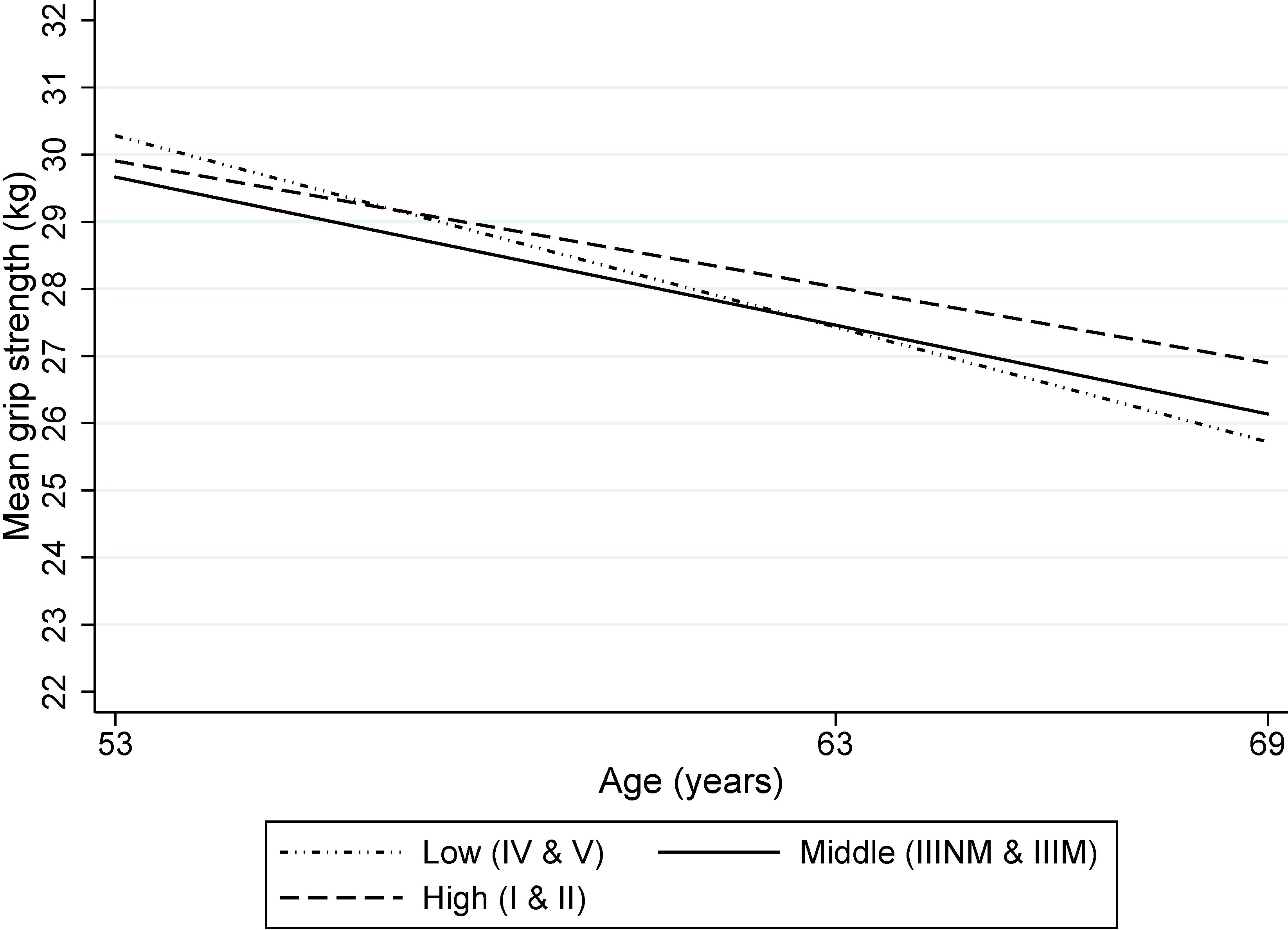
Mean grip strength (kg) by father’s social class for women of mean height (based on 1443 women)

Mutual adjustment of birthweight, physical growth, age at walking (men), age at standing (women), and cognitive development and childhood SEP identified the factors most associated with stronger grip. In men, these were higher birthweight and childhood weight, later puberty, the non-linear relationship with age at first walking and the non-linear relationship with childhood cognition which became increasingly linear with age (Table 3).In women, these were earlier age at first standing, and higher childhood cognition; the estimates for the growth parameters were somewhat reduced in this sample with complete childhood data (Table 4).

**Table 3.**
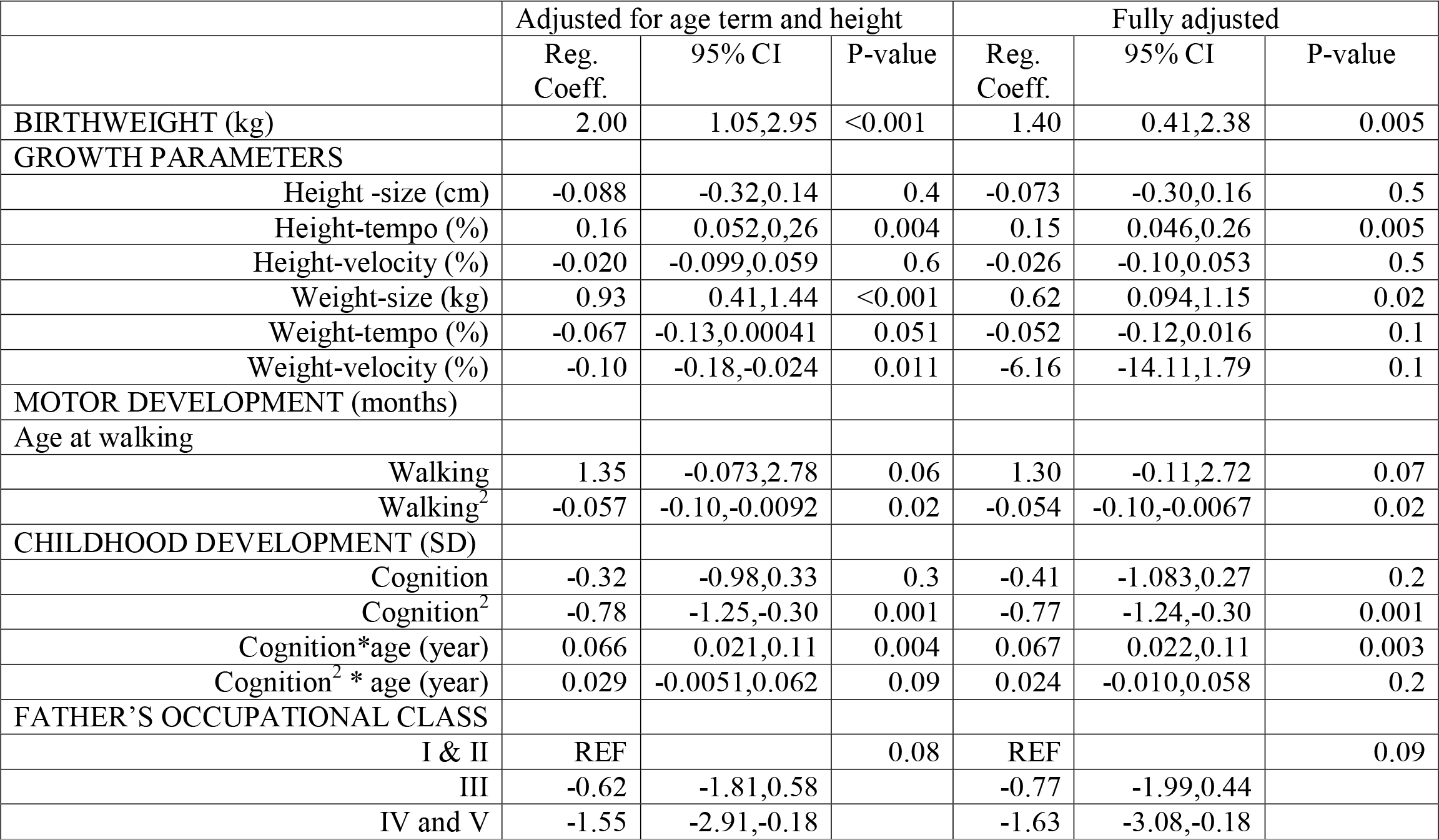
Estimates From Multilevel Models Showing Mean Differences in Grip Strength (kg) and Mean Differences in Grip Strength Change (kg/year) in 1,316 NSHD Men (2,983 Observations) for Mutually Adjusted Childhood Factors. All Models Adjusted for Age Term and Standardised Adult Height.

|  | Adjusted for age term and height |  |  | Fully adjusted |  |  |
| --- | --- | --- | --- | --- | --- | --- |
|  | Reg. Coeff. | 95% CI | P-value | Reg. Coeff. | 95% CI | P-value |
| BIRTHWEIGHT (kg) | 2.00 | 1.05,2.95 | <0.001 | 1.40 | 0.41,2.38 | 0.005 |
| GROWTH PARAMETERS |  |  |  |  |  |  |
| Height -size (cm) | -0.088 | -0.32,0.14 | 0.4 | -0.073 | -0.30,0.16 | 0.5 |
| Height-tempo (%) | 0.16 | 0.052,0.26 | 0.004 | 0.15 | 0.046,0.26 | 0.005 |
| Height-velocity (%) | -0.020 | -0.099,0.059 | 0.6 | -0.026 | -0.10,0.053 | 0.5 |
| Weight-size (kg) | 0.93 | 0.41,1.44 | <0.001 | 0.62 | 0.094,1.15 | 0.02 |
| Weight-tempo (%) | -0.067 | -0.13,0.00041 | 0.051 | -0.052 | -0.12,0.016 | 0.1 |
| Weight-velocity (%) | -0.10 | -0.18,-0.024 | 0.011 | -6.16 | -14.11,1.79 | 0.1 |
| MOTOR DEVELOPMENT (months) |  |  |  |  |  |  |
| Age at walking |  |  |  |  |  |  |
| Walking | 1.35 | -0.073,2.78 | 0.06 | 1.30 | -0.11,2.72 | 0.07 |
| Walking<sup>2</sup> | -0.057 | -0.10,-0.0092 | 0.02 | -0.054 | -0.10,-0.0067 | 0.02 |
| CHILDHOOD DEVELOPMENT (SD) |  |  |  |  |  |  |
| Cognition | -0.32 | -0.98,0.33 | 0.3 | -0.41 | -1.083,0.27 | 0.2 |
| Cognition<sup>2</sup> | -0.78 | -1.25,-0.30 | 0.001 | -0.77 | -1.24,-0.30 | 0.001 |
| Cognition*age (year) | 0.066 | 0.021,0.11 | 0.004 | 0.067 | 0.022,0.11 | 0.003 |
| Cognition<sup>2</sup> * age (year) | 0.029 | -0.0051,0.062 | 0.09 | 0.024 | -0.010,0.058 | 0.2 |
| FATHER'S OCCUPATIONAL CLASS |  |  |  |  |  |  |
| I & II | REF |  | 0.08 | REF |  | 0.09 |
| III | -0.62 | -1.81,0.58 |  | -0.77 | -1.99,0.44 |  |
| IV and V | -1.55 | -2.91,-0.18 |  | -1.63 | -3.08,-0.18 |  |

**Table 4.** Estimates From Multilevel Models Showing Mean Differences in Grip Strength (kg) and Mean Differences in Grip Strength Change (kg/year) in 1,320 NSHD Women (3,069 observations) for Mutually Adjusted Childhood Factors. All Models Adjusted for Age Term and Standardised Adult Height.

|  | Adjusted for age term and height |  |  | Fully adjusted |  |  |
| --- | --- | --- | --- | --- | --- | --- |
|  | Reg. Coeff. | 95% CI | P-value | Reg. Coeff. | 95% CI | P-value |
| BIRTHWEIGHT (kg) | 0.57 | -0.062,1.20 | 0.08 | 0.34 | -0.31,0.98 | 0.3 |
| GROWTH PARAMETERS |  |  |  |  |  |  |
| Height -size (cm) | -0.14 | -0.28,0.0098 | 0.07 | -0.14 | -0.29,-0.0078 | 0.06 |
| Height-tempo (%) | 0.017 | -0.041,0.077 | 0.6 | 0.016 | -0.043,0.075 | 0.6 |
| Height-velocity (%) | -0.0077 | -0.052,0.037 | 0.7 | -0.0056 | -0.054,0.039 | 0.8 |
| Weight-size (kg) | 0.36 | -0.020,0.75 | 0.06 | 0.31 | -0.077,0.71 | 0.1 |
| Weight-tempo (%) | -0.00035 | -0.052,0.052 | 1.0 | 0.008 | -0.042,0.062 | 0.8 |
| Weight-velocity (%) | -0.050 | -0.11,0.0066 | 0.08 | -0.042 | -0.99,0.15 | 0.1 |
| MOTOR DEVELOPMENT (months) |  |  |  |  |  |  |
| Age at standing | -0.16 | 2.50,3.39 | 0.03 | -0.15 | -0.29,-0.0025 | 0.05 |
| CHILDHOOD DEVELOPMENT (SD) |  |  |  |  |  |  |
| Cognition | 0.41 | 0.089,0.73 | .01 | 0.40 | 0.053,0.74 | 0.02 |
| FATHER'S OCCUPATIONAL CLASS |  |  |  |  |  |  |
| I & II | REF |  | 0.2 | REF |  | 0.2 |
| III | -0.59 | -1.65,0.47 |  | -0.45 | -1.53,0.62 |  |
| IV and V | 0.22 | -0.99,1.43 |  | 0.52 | -0.72,1.76 |  |
| III *age (year) | -0.020 | -0.097,0.056 | 0.2 | -0.020 | -0.97,0.056 | 0.1 |
| IV and V*age (year) | -0.079 | -0.17,0.0092 |  | -0.083 | -0.17,0.0056 |  |

Which adult factors account for the associations between developmental factors and grip strength?

All adult covariates were associated with adult grip strength (supplementary Table 2a-g). Of particular relevance for our hypotheses, higher BMI was associated with stronger grip for men (but not women) and this association levelled off at higher levels of BMI and became weaker with age. Having more health conditions was associated with lower grip strength; for men this association strengthened with age but for women it weakened. Higher educational levels were associated with stronger grip, especially among women. The association between verbal memory and grip strength was slightly negative in men and not evident in women at age 53 but grew stronger and positive with age (by 0.10kg/year, 95% CI 0.061,0.15 per 1SD, P <0.001).

In men, the estimates for birthweight, height tempo and weight 0-26 years changed little after adjusting in turn for each adult factor with the greatest reduction in the estimate for birthweight occurring when health conditions were included in the model (Table 5). In contrast, the increasing association between childhood cognitive ability and grip strength with age was strongly attenuated by including verbal memory and a verbal memory by age interaction in the model (Table 6); other adult risk factors had much less impact. The estimate for age at walking was lower in the sample with complete data on adult covariates; however this lower estimate was little affected by adjusting for adult factors (Supplementary Table 3).

**Table 5.**
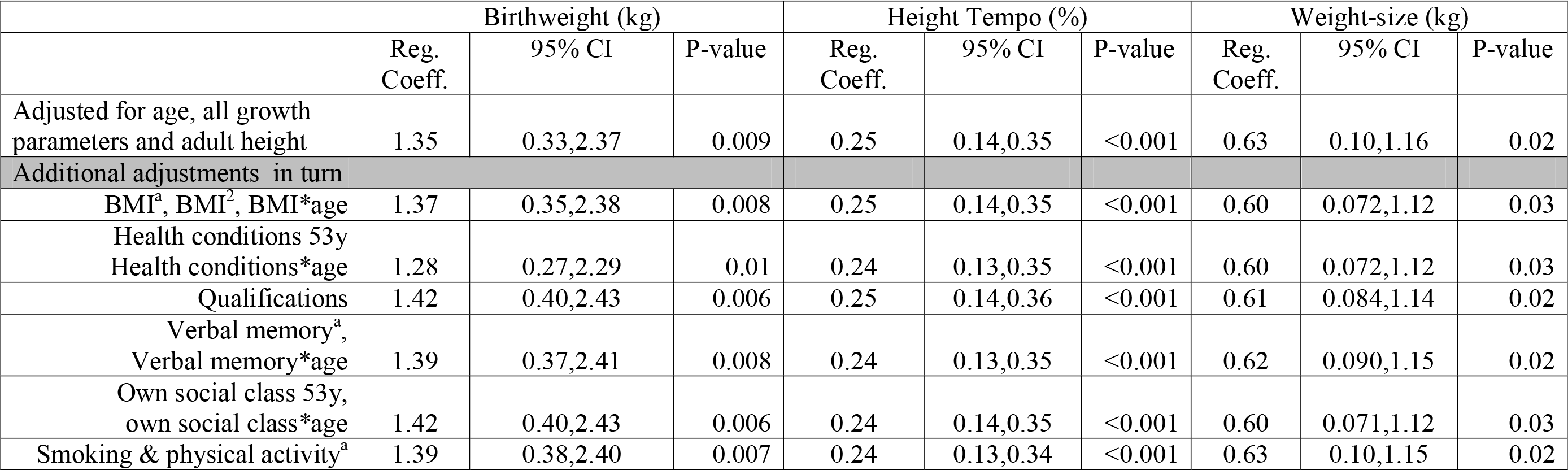
Estimates From Multilevel Models Showing Mean Differences in Grip Strength (kg) by Birthweight, Height Tempo and Weight-Size in 1,295 NSHD Men (2,788 Observations), Adjusted for Age Term, Standardised Adult Height and all Growth Parameters, and Then Additionally Adjusted for Each Set of Adult Factors in Turn

|  | Birthweight (kg) |  |  | Height Tempo (%) |  |  | Weight-size (kg) |  |  |
| --- | --- | --- | --- | --- | --- | --- | --- | --- | --- |
|  | Reg. Coeff. | 95% CI | P-value | Reg. Coeff. | 95% CI | P-value | Reg. Coeff. | 95% CI | P-value |
| Adjusted for age, all growth parameters and adult height | 1.35 | 0.33,2.37 | 0.009 | 0.25 | 0.14,0.35 | <0.001 | 0.63 | 0.10,1.16 | 0.02 |
| Additional adjustments in turn |  |  |  |  |  |  |  |  |  |
| BMI<sup>a</sup>, BMI<sup>2</sup>, BMI*age | 1.37 | 0.35,2.38 | 0.008 | 0.25 | 0.14,0.35 | <0.001 | 0.60 | 0.072,1.12 | 0.03 |
| Health conditions 53y<br>Health conditions*age | 1.28 | 0.27,2.29 | 0.01 | 0.24 | 0.13,0.35 | <0.001 | 0.60 | 0.072,1.12 | 0.03 |
| Qualifications | 1.42 | 0.40,2.43 | 0.006 | 0.25 | 0.14,0.36 | <0.001 | 0.61 | 0.084,1.14 | 0.02 |
| Verbal memory<sup>a</sup>,<br>Verbal memory*age | 1.39 | 0.37,2.41 | 0.008 | 0.24 | 0.13,0.35 | <0.001 | 0.62 | 0.090,1.15 | 0.02 |
| Own social class 53y,<br>own social class*age | 1.42 | 0.40,2.43 | 0.006 | 0.24 | 0.14,0.35 | <0.001 | 0.60 | 0.071,1.12 | 0.03 |
| Smoking & physical activity<sup>a</sup> | 1.39 | 0.38,2.40 | 0.007 | 0.24 | 0.13,0.34 | <0.001 | 0.63 | 0.10,1.15 | 0.02 |

**Table 6.**
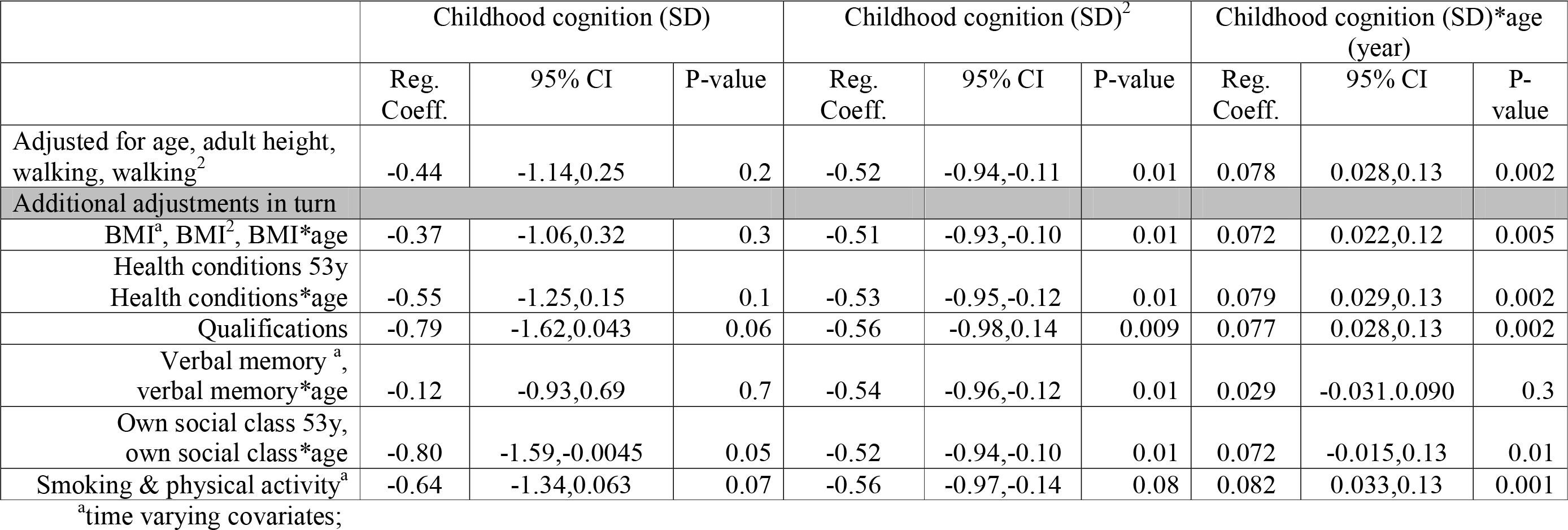
Estimates From Multilevel Models Showing Mean Differences in Grip Strength (kg) and Mean Differences in Grip Strength Change (kg/year) by Childhood Cognition in 1,161 NSHD Men (2,515 Observations), Adjusted for Age Term, Standardised Adult height and Age at First Walking, and Then Additionally Adjusted for Each Set of Adult Factors in Turn

In women, the association between childhood cognitive ability and grip strength was reduced by several adult factors, but especially by educational level (Table 7). However, the inverse association between age at first standing and grip strength in women was not reduced by any of the adult factors.

**Table 7.**
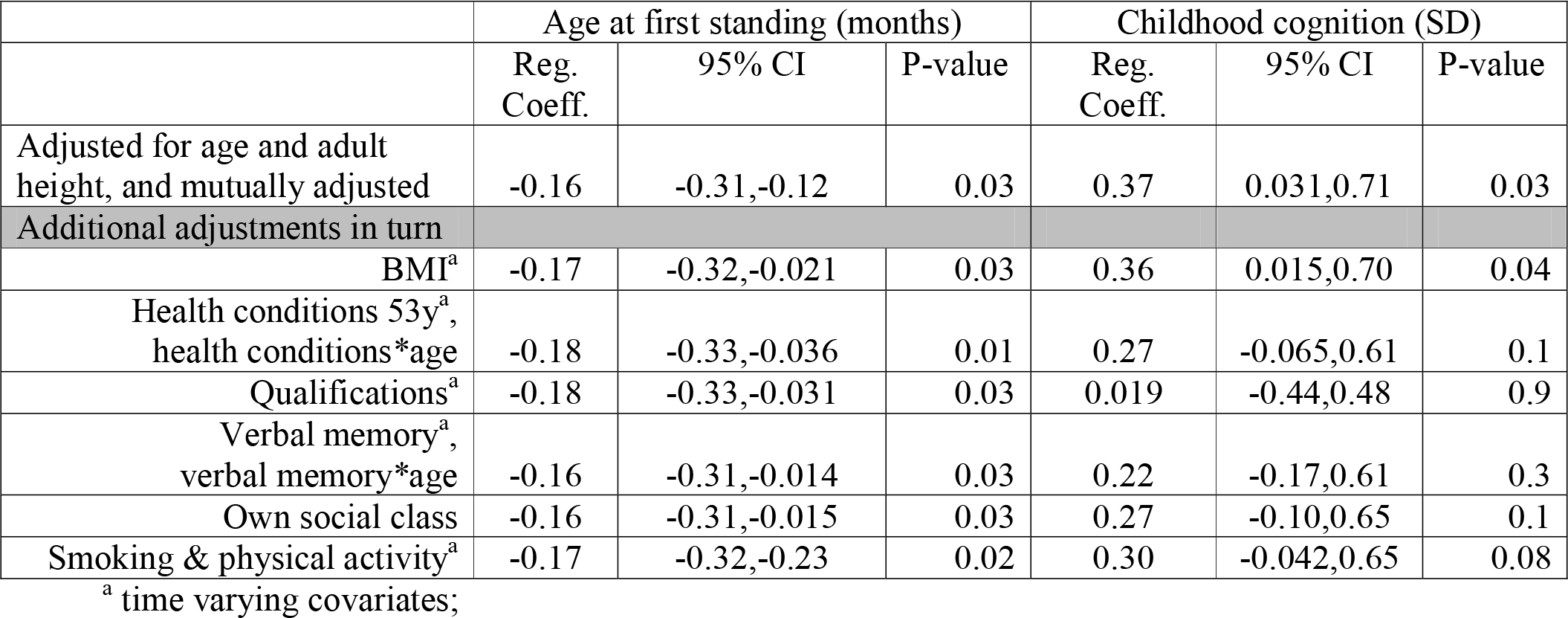
Estimates From Multilevel Models Showing Mean Differences in Grip Strength (kg) by Age at First Standing and Childhood Cognition in 1,211 NSHD Women (2,709 Observations), Adjusted for Age Term and Standardised Adult Height, and Then Additionally Adjusted for Each Set of Adult Factors in Turn.

### Sensitivity analyses

Giving a value for grip strength representing the mid-point of the lowest sex specific fifth (maximum of 81 observations) for those who had no measure because of health reasons had either no change or marginal change on the estimates. Compared with participants who were followed up and assessed at age 69, those who died during follow-up had lower mean grip strength at age 53 (-2.4kg, 95% CI −4.4,-0.41 for men, P=0.02; and −2.5kg, 95% CI −3.9,−1.2 for women, P<0.001) and, for men, there was weak evidence of a faster decline in grip strength (by −0.23kg/year, 95% CI −0.37,0.014, P=0.06). The mean differences in grip strength between those dropping out for reasons other than death and those completing follow-up were smaller (−0.99kg, 95%CI −2.4,0.41 for men, P=0.2; and −1.0kg, 95% CI −1.9, 0.16 for women, P=0.02) with no evidence of age interaction. However, estimates for the associations between developmental and adult risk factors and grip strength remained similar after adjusting for death and attrition.

## DISCUSSION

In a prospective, nationally representative British birth cohort, developmental indicators had persisting associations with grip strength over 16 years from midlife to old age. Physical growth, in terms of heavier birthweight and later puberty and greater weight throughout the growth period was associated with stronger grip in men, and these effects were robust to adjustment for adult factors. ‘On time’ motor development for males and advanced motor development for females were associated with stronger grip which were unexplained by adult factors. Childhood cognitive ability was associated with stronger grip (women) and a slower decline in grip strength (men) during that same life stage; these associations were mediated by later education, or adult cognition.

Comparisons between studies are difficult because different methods have been used to assess decline in grip strength, not all analyses adjust for current adult height, and few studies have developmental data. Some studies have only two grip assessments,(25, 30, 33, 34, 49) or take the difference between the average of several later assessments from the average of several earlier assessments:(32) both methods are limited in their ability to analyse risk factors related to change. Studies with three assessments used bivariate growth curve models(37) or multilevel models.(27) Other studies with five or more assessments used latent growth curve modelling.(28, 50)

In NSHD there was no association between childhood SEP and adult grip strength or its decline after adjusting for developmental factors. Findings from other studies have not been consistent and have adjusted for few other, if any, childhood factors, or have not studied change.(34, 38, 50, 51) The results of a meta-analysis showed modest associations between childhood SEP and adult grip strength at a single time point which were attenuated by adult SEP and current body size, but there was considerable heterogeneity between studies.(52)

The constant effect of birthweight on adult grip strength is consistent with a meta-analysis;(53) this showed a larger estimate for men than women (as this study found) but the sex interaction was not significant. The persistence of the birthweight - grip strength association is worth noting given that more proximal factors may come into play as people age which could have diminished this association. The persisting associations between growth parameters, motor milestones and grip strength are novel findings, and build on previous NSHD work relating to grip strength at age 53,(51) and bone phenotype at age 60-64.(42, 54) Later puberty (in men) was associated with stronger grip, yet earlier puberty was associated with greater areal and volumetric bone mineral density in this cohort,(42) perhaps due to the differential impact of hormonal regulation. Nevertheless, we found that, controlling for contemporaneous body size, greater weight and slower weight velocity throughout the growth period was associated with both greater grip strength and greater bone size,(41) suggesting an extended growth period may benefit both. This could also be the explanation for the persisting associations between motor milestones and grip strength and, in women, for the inverse association between height during growth and later grip strength, after controlling for adult height.

While the associations between physical growth and grip strength were generally independent of adult covariates, the number of health conditions attenuated the birthweight effect more than other covariates. Lower birthweight is predictive of CVD and diabetes,(55) as is poor muscle strength,(16) and may reflect a common pathway to later disease.

The most striking observations in this study were the strengthening of the positive associations between cognition and grip strength with age, whether cognition was assessed in childhood or adult life. This extends an earlier NSHD study showing that the group with meaningful decline in grip strength between ages 53 and 60-64 had lower childhood cognitive ability than those who experienced no meaningful change.(34) In older cohorts, there is growing evidence that changes in grip strength are related to baseline cognition, and that cognitive decline may precede declines in strength,(24) although the few studies investigating covariation in cognition and grip strength have been inconsistent.(37, 56) Our findings complement the findings from older cohorts as they cover midlife changes in grip strength over a longer follow-up period than most previous studies.

A notable implication of our findings regarding lifetime cognition and grip strength is that neural processes have greater impact on grip strength at older ages than in midlife. The attenuation of the childhood cognitive associations once verbal memory or education were taken into account suggests that neurodevelopmental processes play a role in maximising muscle function at maturity but neurodegenerative processes increasingly drive the age-related decline in muscle function. A theoretical model arising out of a review of cognitive ageing, motor learning and motor skills(57) predicts that ageing impairs cognitive functions before affecting the motor systems and that at older ages the connection between cognition and action becomes stronger, as suggested by our findings. To what extent our findings reflect a direct pathway between brain ageing and muscle strength,(58) or shared mechanisms relating, for example, to haemostatic dysregulation or inflammatory processes,(24, 59) is yet to be clarified.

The strengths of NSHD are that it is one of the very few studies with prospectively assessed factors from development onwards, a wide range of potential covariates, and repeat measures of grip strength assessed over a relatively long follow-up period during a critical phase of age-related change. So far these repeat measures cover midlife to early old age, a period which has been studied less often than later ages. NSHD remains broadly representative of the population born in Britain in the early post war period.(60) One limitation is that it is only possible to model linear change as there are currently only three assessments of grip strength. However we did investigate whether each association strengthened or weakened with age. Inevitably there were missing data but neither accounting for deaths and attrition, nor including those unable for health reasons, altered our findings.

In conclusion, patterns of early growth, attainment of motor milestones, and lifetime cognition have persisting associations with grip strength between midlife and old age, even after taking account of adult body size, health conditions and health behaviours. The impact of neural processes strengthened over this stage of life suggesting that at older ages grip strength increasingly reflects both physical and cognitive ageing processes. Interventions that promote muscle development by targeting the developmental factors identified in this study, or maintain peak muscle strength across midlife, by also targeting adult risk factors should increase the chance of an active and independent old age.

## Acknowledgements

We thank NSHD study members for their lifelong participation and past and present members of the NSHD study team who helped to collect the data.

## Abbreviations

BMI: Body mass index
CI: Confidence interval
NSHD: National Survey of Health and Development
SD: Standard deviation
SEP: Socioeconomic position

## Reference List

1. Kuh D, Karunananthan S, Bergman H, Cooper R. A life-course approach to healthy ageing: maintaining physical capability. Proc Nutr Soc. 2014;73(2)237–48.

2. Briggs AM, Cross MJ, Hoy DG, et al. Musculoskeletal Health Conditions Represent a Global Threat to Healthy Aging: A Report for the 2015 World Health Organization World Report on Ageing and Health. Gerontologist. 2016;56 Suppl 2:S243–55.

3. Wolfe RR. The underappreciated role of muscle in health and disease. Am J Clin Nutr. 2006;84(3)475–82.

4. Manring H, Abreu E, Brotto L, et al. Novel excitation-contraction coupling related genes reveal aspects of muscle weakness beyond atrophy-new hopes for treatment of musculoskeletal diseases. Frontiers Physiol. 2014;5:37.

5. Bohannon RW. Are hand-grip and knee extension strength reflective of a common construct? Percept Mot Skills. 2012;114(2)514–8.

6. Martin-Ruiz CM, von Zglinicki T. A life course approach to biomarkers of ageing. In: Kuh D, Cooper R, Hardy R, Richards M, Ben-Shlomo Y, editors. A life course approach to healthy ageing. 1st ed. Oxford: Oxford University Press; 2014. p. 177–86.

7. Sayer AA, Kirkwood TB. Grip strength and mortality: a biomarker of ageing? Lancet. 2015;386(9990)226–7.

8. Justice JN, Cesari M, Seals DR, et al. Comparative Approaches to Understanding the Relation Between Aging and Physical Function. J Gerontol A Biol Sci Med Sci. 2016;71(10) 1243–53.

9. Dodds RM, Syddall HE, Cooper R, et al. Grip strength across the life course: normative data from twelve British studies. PLoS ONE. 2014;9(12)e113637.

10. Dodds RM, Syddall HE, Cooper R, et al. Global variation in grip strength: a systematic review and meta-analysis of normative data. Age Ageing. 2016;45(2)209–16.

11. Nahhas RW, Choh AC, Lee M, et al. Bayesian longitudinal plateau model of adult grip strength. Am J Hum Biol. 2010;22(5)648–56.

12. Rantanen T, Volpato S, Ferrucci L, et al. Handgrip strength and cause-specific and total mortality in older disabled women: exploring the mechanism. J Am.Geriatr Soc. 2003;51(5)636–41.

13. Cooper R, Kuh D, Hardy R. Objectively measured physical capability levels and mortality: systematic review and meta-analysis. BMJ 2010;341:c4467.

14. Cooper R, Kuh D, Cooper C, et al. Objective measures of physical capability and subsequent health: a systematic review. Age Ageing. 2011;40:14–23.

15. den Ouden ME, Schuurmans MJ, Arts IE, van der Schouw YT. Physical performance characteristics related to disability in older persons: a systematic review. Maturitas. 2011;69(3)208–19.

16. Leong DP, Teo KK, Rangarajan S, et al. Prognostic value of grip strength: findings from the Prospective Urban Rural Epidemiology (PURE) study. Lancet. 2015;386(9990)266–73.

17. Ortega FB, Silventoinen K, Tynelius P, Rasmussen F. Muscular strength in male adolescents and premature death: cohort study of one million participants. BMJ. 2012;345:e7279.

18. Kim Y, Wijndaele K, Lee DC, et al. Independent and joint associations of grip strength and adiposity with all-cause and cardiovascular disease mortality in 403,199 adults: the UK Biobank study. Am J Clin Nutr. 2017;106(3)773–82.

19. Peterson MD, Zhang P, Duchowny KA, et al. Declines in Strength and Mortality Risk Among Older Mexican Americans: Joint Modeling of Survival and Longitudinal Data. J Gerontol A Biol Sci Med Sci. 2016;71(12)1646–52. doi:10.1093/gerona/glw051

20. Syddall HE, Westbury LD, Dodds R, et al. Mortality in the Hertfordshire Ageing Study: association with level and loss of hand grip strength in later life. Age Ageing. 2017;46(3)407–12.

21. Shaw SC, Dennison EM, Cooper C. Epidemiology of Sarcopenia: Determinants Throughout the Lifecourse. Calcif.Tissue Int. 2017;101(3)229–47.

22. Clouston SAP, Brewster P, Kuh D, et al. The dynamic relationship between physical function and cognition in longitudinal aging cohorts. Epidemiologic Reviews. 2013;35(1)33–50.

23. Syddall HE, Westbury LD, Shaw SC, et al. Correlates of Level and Loss of Grip Strength in Later Life: Findings from the English Longitudinal Study of Ageing and the Hertfordshire Cohort Study. Calcif Tissue Int. 2017.

24. Fritz NE, McCarthy CJ, Adamo DE. Handgrip strength as a means of monitoring progression of cognitive decline - A scoping review. Ageing Res Rev. 2017;35:112–23.

25. Cooper R, Muniz-Terrera G, Kuh D. Associations of behavioural risk factors and health status with changes in physical capability over 10 years of follow-up: the MRC National Survey of Health and Development. BMJ Open. 2016;6(4)e009962.

26. Granic A, Davies K, Jagger C, et al. Grip Strength Decline and Its Determinants in the Very Old: Longitudinal Findings from the Newcastle 85+ Study. PLoS.One. 2016;11(9)e0163183.

27. Botoseneanu A, Bennett JM, Nyquist L, et al. Cardiometabolic Risk, Socio-Psychological Factors, and Trajectory of Grip Strength Among Older Japanese Adults. J Aging Health. 2015;27(7)1123–46.

28. Sternang O, Reynolds CA, Finkel D, et al. Factors associated with grip strength decline in older adults. Age Ageing. 2015;44(2)269–74.

29. Stenholm S, Tiainen K, Rantanen T, et al. Long-Term Determinants of Muscle Strength Decline: Prospective Evidence from the 22-Year Mini-Finland Follow-Up Survey. J Am Geriat Soc. 2012;60(1)77–85.

30. Miller DK, Malmstrom TK, Miller JP, et al. Predictors of change in grip strength over 3 years in the African American health project. J.Aging Health. 2010;22(2)183–96.

31. Rantanen T, Penninx BW, Masaki K, Lintunen T, Foley D, Guralnik JM. Depressed mood and body mass index as predictors of muscle strength decline in old men. J Am Geriatr Soc. 2000;48(6)613–7.

32. Rantanen T, Masaki K, Foley D, et al. Grip strength changes over 27 yr in Japanese-American men. J App Physiol. 1998;85(6)2047–53.

33. Clement FJ. Longitudinal and cross-sectional assessments of age changes in physical strength as related to sex, social class, and mental ability. J Gerontol. 1974;29:423–9.

34. Cooper R, Richards M, Kuh D. Childhood Cognitive Ability and Age-Related Changes in Physical Capability From Midlife: Findings From a British Birth Cohort Study. Psychosom Med. 2017;79(7)785–91.

35. Aihie Sayer A, Cooper C, Evans JR, et al. Are rates of ageing determined in utero? Age Ageing. 1998;27:579–83.

36. Cooper R, Hardy R, Sayers A, Kuh D. A life course approach to physical capability. In: Kuh D, Cooper R, Hardy R, Richards M, Ben-Shlomo Y, editors. A life course approach to healthy ageing. 1st ed. Oxford: Oxford University Press; 2014. p. 16–31.

37. Deary IJ, Johnson W, Gow AJ, et al. Losing one’s grip: a bivariate growth curve model of grip strength and nonverbal reasoning from age 79 to 87 years in the Lothian Birth Cohort 1921. J Gerontol B Psychol Sci Soc Sci. 2011;66(6)699–707.

38. Starr JM, Deary IJ. Socio-economic position predicts grip strength and its decline between 79 and 87 years: the Lothian Birth Cohort 1921. Age and Ageing. 2011;40(6)749–52.

39. Kuh D, Wong A, Shah I, et al. The MRC National Survey of Health and Development reaches age 70: maintaining participation at older ages in a birth cohort study. Eur J Epidemiol. 2016;31(11) 1135–47.

40. Lessof C, Wong A, Bendayan R et al. Testing for differences in measurement devices: findings from a randomized trial to compare measures of physiological function and physical performance European Survey Research Association; 2017 19th July; Lisbon.

41. Cole TJ, Kuh D, Johnson W, et al. Using Super-Imposition by Translation And Rotation (SITAR) to relate pubertal growth to bone health in later life: the Medical Research Council (MRC) National Survey of Health and Development. Int J Epidemiol. 2016;45(4) 1125–34.

42. Kuh D, Muthuri SG, Moore A, et al. Pubertal timing and bone phenotype in early old age: findings from a British birth cohort study. Int J Epidemiol. 2016;45(4)1113–24.

43. Heim AW The AH4 group test of intelligence. Windsor: NFER-Nelson; 1970.

44. Pigeon DA. Details of the fifteen years tests. In: JWB D, JM R, HR S, editors. All our future. London: Davies; 1968. p. Appendix 1.

45. Davis D, Bendayan R, Muniz G, et al. Decline in search speed and verbal memory over 26 years of midlife in a British birth cohort. Neuroepidemiology. 2017;. 2017;49(3-4):121–128.

46. Oksuzyan A, Maier H, McGue M, et al. Sex Differences in the Level and Rate of Change of Physical Function and Grip Strength in the Danish 1905-Cohort Study. J Aging Health. 2010;22(5)589–610.

47. Botoseneanu A, Allore HG, Mendes de Leon CF, Gahbauer EA, Gill TM. Sex Differences in Concomitant Trajectories of Self-Reported Disability and Measured Physical Capacity in Older Adults. J Gerontol A Biol Sci Med Sci. 2016;71(8)1056–62.

48. Botoseneanu A, Liang J. The effect of stability and change in health behaviors on trajectories of body mass index in older Americans: a 14-year longitudinal study. J Gerontol A Biol Sci Med Sci. 2012;67(10)1075–84. doi:10.1093/gerona/gls073

49. Daly RM, Ahlborg HG, Ringsberg K, et al. Association between changes in habitual physical activity and changes in bone density, muscle strength, and functional performance in elderly men and women. J Am Geriatr Soc. 2008;56(12)2252–60.

50. Kroger H, Fritzell J, Hoffmann R. The Association of Levels of and Decline in Grip Strength in Old Age with Trajectories of Life Course Occupational Position. PLoS.One. 2016;11(5)e0155954.

51. Kuh D, Hardy R, Butterworth S, et al. Developmental origins of midlife grip strength: Findings from a birth cohort study. J Gerontol A Biol Sci Med Sci. 2006;61(7)702–6.

52. Birnie K, Cooper R, Martin RM, et al. Childhood socioeconomic position and objectively measured physical capability levels in adulthood: A systematic review and metaanalysis. PLoS ONE. 2011;6(1)e15564.

53. Dodds R, Denison HJ, Ntani G, et al. Birth weight and muscle strength: A systematic review and meta-analysis. J Nutr Health Ageing. 2012;16(7)609–15.

54. Kuh D, Wills AK, Shah I, et al. Growth from birth to adulthood and bone phenotype in early old age: A British birth cohort study. J Bone Miner Res. 2014;29(1)123–33.

55. Whincup PH, Kaye SJ, Owen CG, et al. Birth weight and risk of type 2 diabetes: A systematic review. JAMA. 2008;300(24)2886–97.

56. Praetorius Bjork M, Johansson B, Hassing LB. I forgot when I lost my grip-strong associations between cognition and grip strength in level of performance and change across time in relation to impending death. Neurobiol Aging. 2016;38:68–72.

57. Ren J, Wu YD, Chan JS, Yan JH. Cognitive aging affects motor performance and learning. Geriat Gerontol Int. 2013;13(1)19–27.

58. L Bhanushali M, Conwit R, Metter RM, Ferrucci L. The role of the nervous system in muscle atrophy. In: Cruz-Jentofy AJ, Morley JE editors. Sarcopenia. Chichester, UK: John Wiley & Sons; 2012.

59. Newman AB, Sanders JL, Kizer JR, et al. Trajectories of function and biomarkers with age: the CHS All Stars Study. Int.J.Epidemiol. 2016;45(4)1135–45.

60. Stafford M, Black S, Shah I, et al. Using a birth cohort to study ageing: representativeness and response rates in the National Survey of Health and Development. Eur J Ageing. 2013;10(2)145–57.

